# CagY-dependent regulation of type IV secretion in *Helicobacter pylori* is associated with alterations in integrin binding

**DOI:** 10.1101/294769

**Authors:** Emma C. Skoog, Vasilios A. Morikis, Miriam E. Martin, Greg A. Foster, Lucy P. Cai, Lori M. Hansen, Beibei Li, Jennifer A. Gaddy, Scott I. Simon, Jay V. Solnick

**Affiliations:** Department of Medicine; Department of Microbiology & Immunology; Department of Biomedical Engineering; Department of Microbiology and Molecular Genetics; Center for Comparative Medicine University of California, Davis School of Medicine Davis, CA 95616 USA; Shanghai Veterinary Research Institute Chinese Academy of Agricultural Science Shanghai 200241, P. R. China; Department of Veterans Affairs Tennessee Valley Healthcare Systems Department of Medicine Vanderbilt University Medical Center Nashville, TN 37212 USA

## Abstract

Strains of *Helicobacter pylori* that cause ulcer or gastric cancer typically express a type IV secretion system (T4SS) encoded by the *cag* pathogenicity island (PAI). CagY is an ortholog of VirB10 that, unlike other VirB10 orthologs, has a large middle repeat region (MRR) with extensive repetitive sequence motifs, which undergo CD4+ T cell-dependent recombination during infection of mice. Recombination in the CagY MRR reduces T4SS function, diminishes the host inflammatory response, and enables the bacteria to colonize at a higher density. Since CagY is known to bind human α_5_β_1_ integrin, we tested the hypothesis that recombination in the CagY MRR regulates T4SS function by modulating binding to α_5_β_1_ integrin. Using a cell-free microfluidic assay, we found that *H. pylori* binding to α_5_β_1_ integrin under shear flow is dependent on the CagY MRR, but independent of the presence of the T4SS pili, which are only formed when *H. pylori* is in contact with host cells. Similarly, expression of CagY in the absence of other T4SS genes was necessary and sufficient for whole bacterial cell binding to α_5_β_1_ integrin. Bacteria with variant *cagY* alleles that reduced T4SS function showed comparable reduction in binding to α_5_β_1_ integrin, though CagY was still expressed on the bacterial surface. We speculate that *cagY-*dependent modulation of *H. pylori* T4SS function is mediated by alterations in binding to α_5_β_1_ integrin, which in turn regulates the host inflammatory response so as to maximize persistent infection.

**IMPORTANCE:** Infection with *H. pylori* can cause peptic ulcers, and is the most important risk factor for gastric cancer, the third most common cause of cancer death worldwide. The major *H. pylori* virulence factor that determines whether infection causes disease or asymptomatic colonization is the type IV secretion system (T4SS), a sort of molecular syringe that injects bacterial products into gastric epithelial cells and alters host cell physiology. We previously showed that recombination in CagY, an essential T4SS component, modulates the function of the T4SS. Here we found that these recombination events produce parallel changes in specific binding to α_5_β_1_ integrin, a host cell receptor that is essential for T4SS-dependent translocation of bacterial effectors. We propose that CagY-dependent binding to α_5_β_1_ integrin acts like a molecular rheostat that alters T4SS function and modulates the host immune response to promote persistent infection.

## INTRODUCTION

*Helicobacter pylori* infection most often causes only asymptomatic gastritis, but is considered an important human pathogen because it is the major risk factor for development of peptic ulcer disease and gastric adenocarcinoma (1), the third most common cause of cancer death. On the other hand, *H. pylori* infection may also have beneficial effects, particularly prevention of chronic diseases that have increased in frequency in developed countries as the prevalence of *H. pylori* has declined (2). The bacterial virulence factor most strongly associated with the outcome of *H. pylori* infection is the *cag* pathogenicity island (*cag*PAI), a ~40 kb DNA segment that encodes a type IV secretion system (T4SS). When *H. pylori* comes in contact with the gastric epithelium, it assembles the T4SS pilus (3), through which it injects the CagA oncoprotein into host cells (4). Other T4SS-dependent effectors have also been identified, including DNA (5), peptidoglycan (6) and heptose-1,7-bisphosphate, a metabolic precursor in lipopolysaccharide biosynthesis (7-9). Together, T4SS injection of effector molecules results in complex changes in host cell physiology that include cytoskeletal rearrangements, disruption of tight junctions, loss in cell polarity, and production of interleukin 8 (IL-8) and other proinflammatory cytokines (4, 10).

Host cell expression of β1 integrins is required for T4SS-dependent translocation of CagA (11, 12), and presumably other effectors as well. Four *cag*PAI proteins essential for T4SS function have been found to bind β_1_ integrins, though the details are unclear and some reports are contradictory. The first to be described was CagL, an RGD-dependent ligand for α_5_β_1_ integrin that presumably mimics fibronectin, an intrinsic host integrin ligand (11). An RGD helper motif in CagL (FEANE) may also be important (13). However, other studies have failed to demonstrate CagL binding to β_1_ integrins (12), have yielded discrepant results about the role of CagL polymorphisms (14-16), or have identified completely different integrin binding partners, including α_V_β_6_ and α_V_β_8_ (17). CagA, CagI and CagY have also been shown to bind β_1_ integrin using yeast two hybrid, immunoprecipitation, and flow cytometry approaches (12). However, *H. pylori* binding to integrins has only occasionally been performed with intact bacterial cells (12, 18), and the role of the *cag*PAI-encoded proteins for integrin binding has not yet been examined in the context of a fully assembled T4SS.

It has long been known that passage of *H. pylori* in mice results in loss of T4SS function (19, 20). We previously demonstrated that this is typically a result of recombination events in *cagY* (21), a *virB10* ortholog that contains in its middle repeat region (MRR) an extraordinary series of direct DNA repeats that are predicted to encode in-frame insertions or deletions in a surface-exposed region of the protein (22). Recombination events in the *cagY* MRR lead to expression of an alternative CagY allele that can modulate T4SS function, including induction of IL-8 and translocation of CagA (21). This modulation can occur in a graded fashion, and cause both gain and loss of T4SS function (21). More recently, we demonstrated that IFNγ and CD4+ T cells are essential for *cagY*-mediated loss of T4SS function, which can rescue colonization in *IL10-/-* mice that have an exaggerated inflammatory response to *H. pylori* infection (23). Together, these results suggest that *cagY* recombination serves as an immune sensitive molecular rheostat that “tunes” the host inflammatory response so as to maintain persistent infection.

Here we examined the mechanism by which recombination in *cagY* alters T4SS function. Since CagY forms the spokes of a T4SS core complex, together with CagX, CagM, CagT and Cag3 (24, 25), one possibility is that changes in the MRR alter T4SS function by modifying essential protein-protein interactions, or changing the pore through which effectors must travel. Alternatively, since CagY recombination occurs in the MRR, which is predicted to extend extracellularly, allelic variation in CagY might alter integrin binding. At first glance this seemed unlikely since there are multiple *cag*PAI proteins that bind integrins. Surprisingly, our results demonstrate that indeed recombination in the CagY MRR alters binding to β_1_ integrin, which in turn modulates T4SS function. Moreover, the CagY MRR is expressed on the bacterial surface even in the absence of a T4SS pilus. We propose that CagY is a bifunctional protein that contains a VirB10 domain that is an essential part of a complete T4SS structure, and an MRR region that mediates close contact to the host cell and modulates T4SS function.

## RESULTS

### *H. pylori* binds to α5β1 integrin in a host cell-free assay

Previous studies analyzed binding of *H. pylori* to β_1_ integrins by protein-protein interaction assays, protein to host cell binding, or bacterial co-localization to β_1_ integrin on host cells *in vitro* (11, 12). To demonstrate binding of intact live *H. pylori* to β_1_ integrin, we developed a microfluidic assay in which human recombinant α_5_β_1_ integrin was coated on glass cover slips, which served as the substrate of a flow channel (Fig 1A). Fluorescently stained bacteria were flowed through the channel at a defined shear stress (~1 dyne/cm^2^), microscopic images were recorded, and immobilized fluorescent bacteria were counted. To validate the microfluidic assay, we first analyzed binding of *Escherichia coli* expressing *Yersinia* invasin, a well-characterized β_1_ integrin ligand (26). *Yersinia* InvA was expressed in *E. coli* after IPTG stimulation (Fig S1A) and was presented on the bacterial cell surface (Fig 1B). *E. coli* harboring the plasmid vector alone served as a negative control. *E. coli* expressing InvA and fluorescently stained with either DiO or DiD showed markedly increased binding to α_5_β_1_ compared to control *E. coli* with vector alone (Fig 1C). Similar results were obtained when InvA and control strains were mixed 1:1 and analyzed simultaneously, which permitted direct comparison in the same flow channel, and limited variability that might otherwise arise from differences in integrin density or flow disturbances on glass coverslips (Fig S1B).

**Fig 1:**
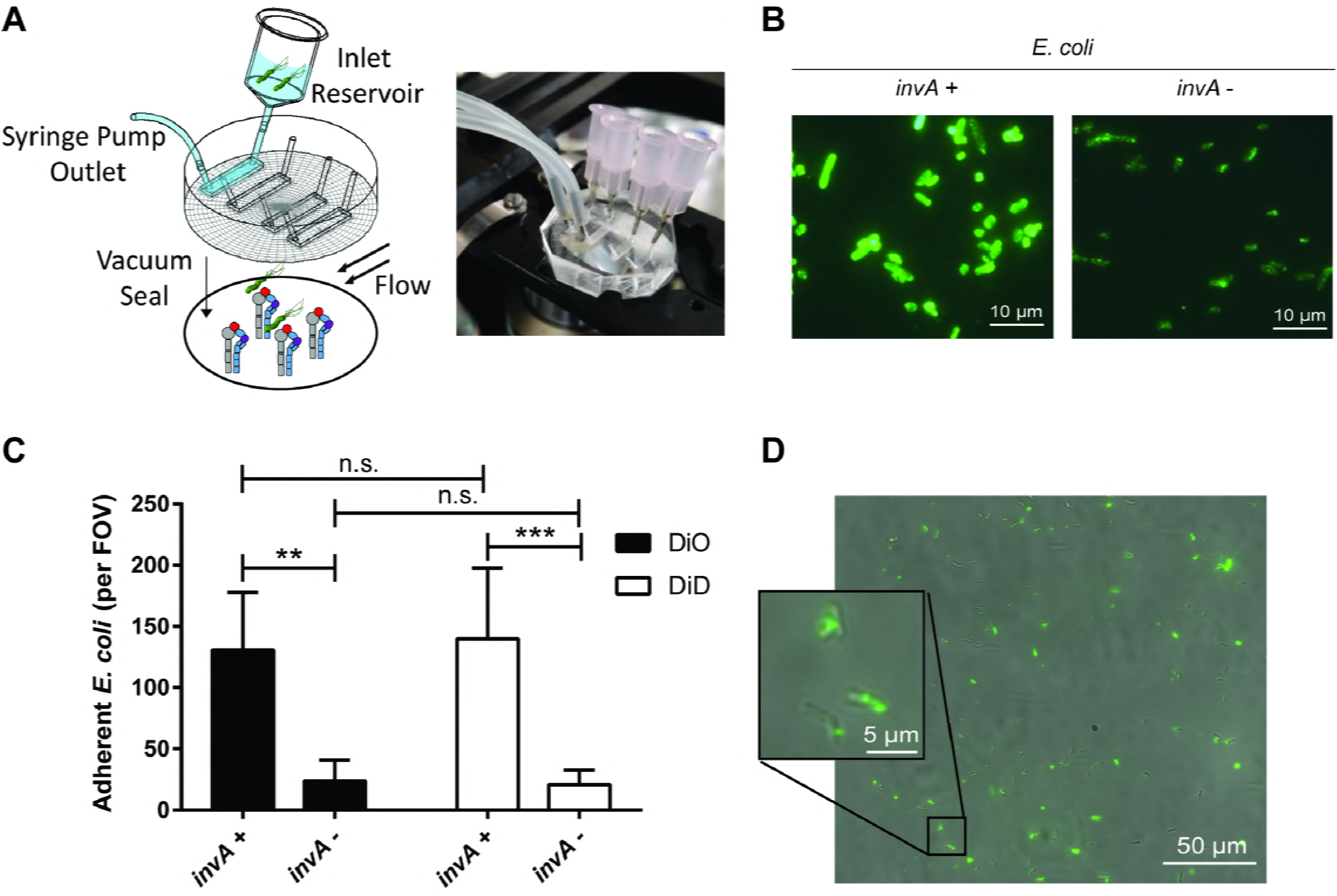
Microfluidic detection of bacterial adherence to recombinant α_5_β_1_ integrin. (A) Schematic diagram and photograph of the microfluidic flow cell assembly. (B) Immunofluorescent detection of InvA on the surface of non-permeabilized IPTG-treated *E. coli* containing the pRI253 plasmid with or without *invA*. (C) Attachment to α_5_β_1_ integrin of IPTG-treated *E. coli*. Each strain was used at an OD_600_ of 0.8, labeled with DiD or DiO and assayed separately. (D) Micrograph of *H. pylori* J166 labeled with DiO membrane dye, attached to α_5_β_1_ integrin in the microfluidic flow cell. Brightfield and fluorescence overlay of the field of view (FOV) demonstrates fluorescent labeling of *H. pylori*.

Fluorescently stained *H. pylori* was also readily visualized adherent to α_5_β_1_ integrin (Fig 1D). *H. pylori* strains J166 and PMSS1 both attached to α_5_β_1_ integrin in a concentration-dependent manner and reached saturation at an optical density (OD_600_) of 0.8 (Fig 2A). This correlates with approximately 4×10^8^ bacterial cells per ml and was used for all subsequent experiments. Binding was blocked by pre-incubating the integrin-coated cover slips with P5D2 anti-β_1_ antibody, which sterically inhibits integrin-dependent binding (Fig 2B). Allosterically stabilizing the α_5_β_1_ integrin in the low affinity conformation by pre-incubation with SG19 antibody decreased *H. pylori*-integrin binding, while the TS2/16 antibody that locks α_5_β_1_ in the high affinity conformation yielded binding similar to that of an isotype control, indicating that the majority of derivatized α_5_β_1_ is active (Fig 2B). This was also supported by the observation that pretreatment of the microfluidic channels with Mn^2+^ to lock α_5_β_1_ integrin in the high affinity state yielded *H. pylori* adherence similar to that with the TS2/16 antibody and the isotype control (Fig 2B). Adherence to α_5_β_1_ integrin was greater than to α_4_β_1_ and α_L_β_2_ (Fig 2C). Thus, live whole cell *H. pylori* binds specifically and in a conformation dependent manner to α_5_β_1_ integrin.

**Fig 2:**
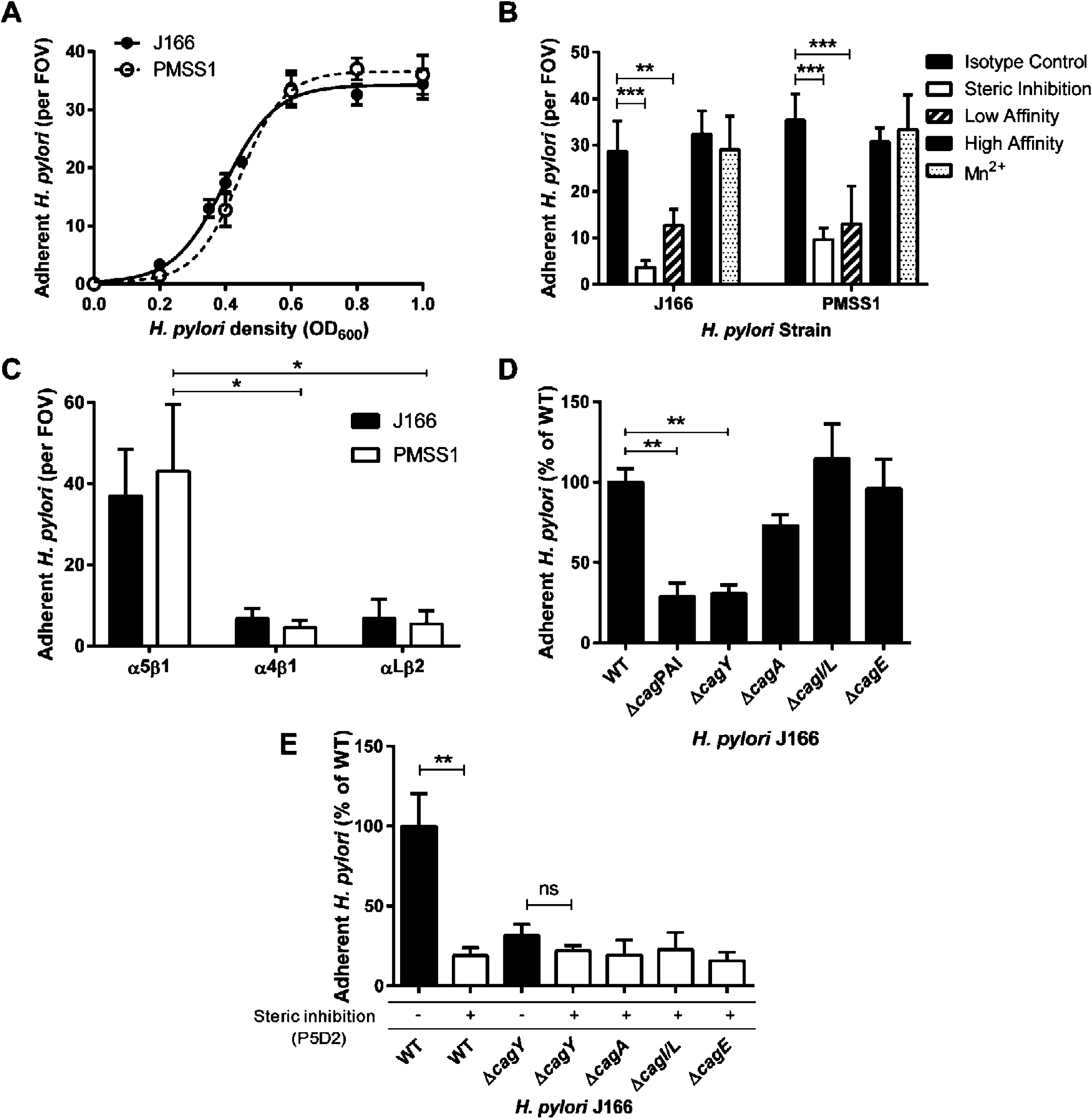
α_5_β_1_ integrin adherence of WT *H. pylori* J166 and *cag*PAI deletion mutants. (A) Adherent *H. pylori* J166 and PMSS1 per FOV as a function of bacterial optical cell density at 600 nm (OD_600_). (B) Adherent *H. pylori* after pre-incubation of flow cells with B11/6 isotype control antibody, P5D2 antibody to sterically inhibit β_1_ integrin binding, or antibodies to lock the integrin in the low (SG19) or high (TS2/16) affinity conformation, respectively. Treatment of integrin with Mn^2+^ to stabilize the high affinity state produced results similar to treatment with TS2/16 and the B11/6 isotype control antibody. (C) Adherence to α_5_β_1_, α_4_β_1_ and α_L_β_2_ integrins. (D) Adherence to α_5_β_1_ integrin of J166 WT and deletion mutants, which were fluorescently labeled with DiO and DiI, respectively, mixed in a 1:1 ratio, and enumerated by counting fluorescent bacteria per field of view (FOV). Results are expressed as the ratio of deletion mutant to WT. (E) Steric inhibition with P5D2 antibody (white bars) demonstrated that adherence is integrin-specific. Δ*cagY* adherence was similar with and without steric inhibition, suggesting that it represents only non-specific background binding. Results are mean (± SEM) of 3 to 5 independent experiments. *P<0.05, **P<0.01, ***P<0.001.

### *H. pylori* adherence to α_5_β_1_ integrin in a host cell-free assay is dependent on CagY

To determine if the *cag*PAI, or any of the putative integrin binding partners (CagA, CagI, CagL or CagY), are responsible for α_5_β_1_ integrin binding of intact *H. pylori*, we compared deletion mutants of *H. pylori* J166 to the wild type (WT) control. The number of adherent mutant and WT *H. pylori* per field of view was determined, and the results were analyzed as the percent adherence of the mutant compared to WT. Initial control experiments demonstrated that WT and selected mutant strains stained with similar efficiency with both dyes (Fig S2A), and that adhesion was independent of the dye and was similar whether strains were analyzed individually or competitively (Fig S2B). Adherence to α_5_β_1_ integrin was markedly reduced by deletion of the entire *cag*PAI (Δ*cag*PAI), but not by deletion of *cagE* (Δ*cagE*) or *cagI/L* (Δ *cagI/L*) (Fig 2D). Deletion of *cagA* (Δ*cagA*) produced a small reduction in adherence to α_5_β_1_ integrin but the difference was not statistically significant (P=0.25). In contrast, integrin adherence by the *cagY* deletion mutant (Δ*cagY*, shown schematically in Fig 3) was significantly reduced to a level similar to Δ*cag*PAI (Fig 2D). Blocking by treatment with anti-β_1_ antibody demonstrated β_1_-specific binding in Δ*cagA*, Δ*cagI/L, and* Δ*cagE* mutants (Fig 2E), which all produced CagY as demonstrated by immunoblot (Fig S3A). Δ*cagY* showed only residual adherence that was not β_1_-specific (Fig 2E). Together, these results demonstrate that in this host cell-free system, adhesion of *H. pylori* to α_5_β_1_ integrin under physiological levels of shear stress is mediated predominantly by CagY.

**Fig 3:**
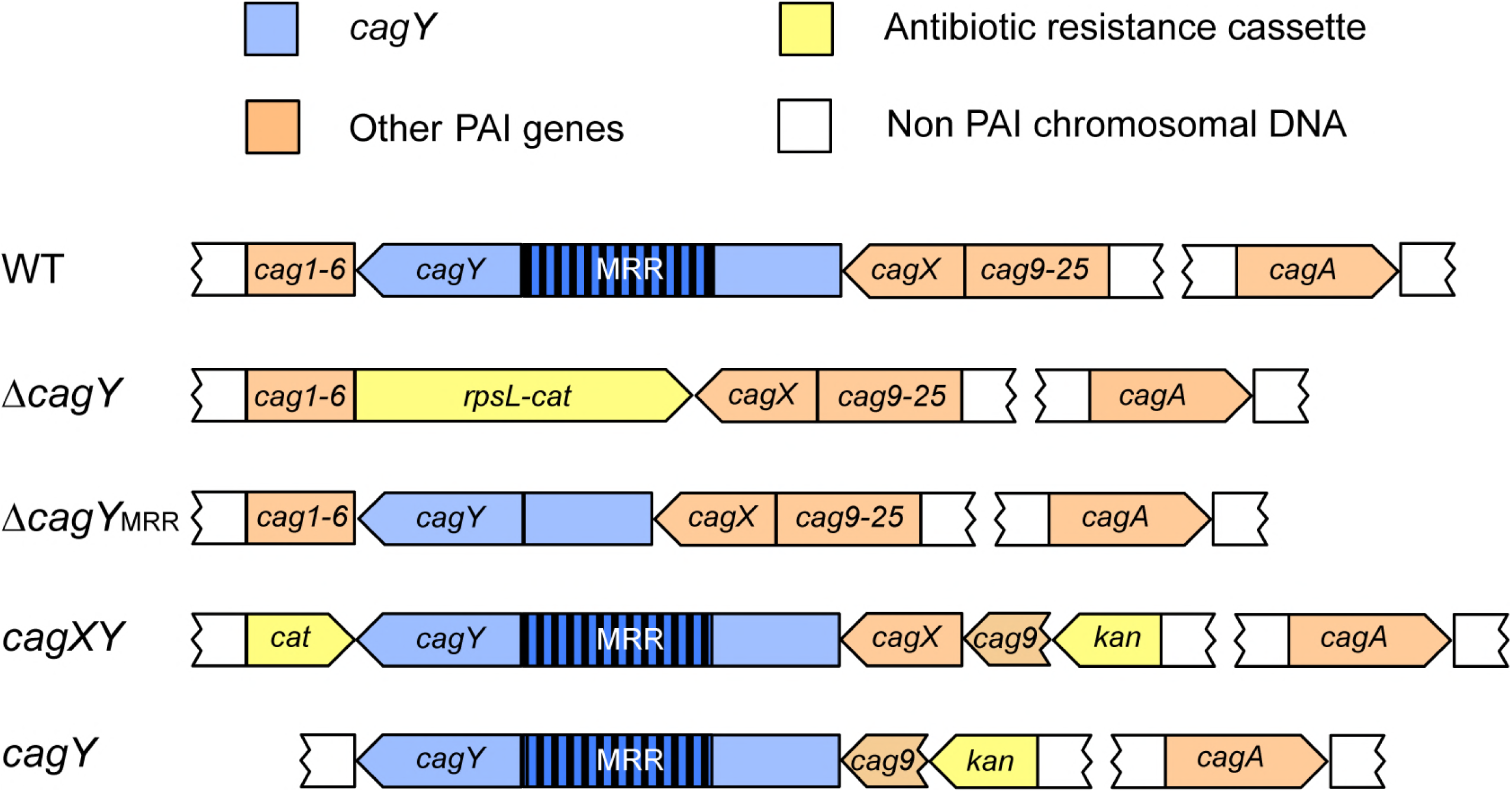
Schematic diagram of the *H. pylori* J166 *cag*PAI in the WT and selected deletion mutants. In J166Δ*cagY*, the entire *cagY* gene is replaced by a *cat-rpsL* cassette (streptomycin susceptibility and chloramphenicol resistance). Δ*cagY*_MRR_ has an unmarked, in-frame deletion of the MRR created by contraselection. In J166 *cagXY*, *cag1-6* is replaced with *cat*, and *cag9-25* is replaced with a kanamycin resistance cassette, starting from after the putative *cagY* promoter in *cag9*. J166 *cagY* has an unmarked deletion of *cag1-6* and *cagX*, while *cag9-25* downstream of the *cagX/Y* promoter in cag9 are replaced with a kanamycin resistance cassette. *cagA* is intact in all strains since it is not on the *cag*PAI in J166 (56).

### CagY-mediated integrin binding is independent of the T4SS pilus

*H. pylori* T4SS pilus formation is thought to require host cell contact (27), though this has never been formally demonstrated. Since *H. pylori* attachment to integrin in the flow channel occurs in the absence of host cells, this suggests that *H. pylori* can bind to α_5_β_1_ integrin independent of the T4SS pilus. To examine this further, we used field emission scanning electron microscopy (FEG-SEM) to image the T4SS pili in *H. pylori* WT and Δ*cag*PAI, co-cultured with or without AGS gastric epithelial cells. Numerous pili were observed on WT *H. pylori* J166, but only in the presence of AGS cells (Fig 4). As expected, no pili were detected on J166Δ*cag*PAI. The same results were found for *H. pylori* PMSS1 WT and Δ*cag*PAI (Fig S4). Culture of *H. pylori* together with α_5_β_1_ integrin also failed to induce pilus formation (data not shown). Therefore, under shear flow in this cell-free system, CagY-mediated binding to α_5_β_1_ integrin does not require formation of the T4SS pilus. To further demonstrate that CagY is sufficient for integrin binding in the absence of the T4SS pilus, all of the PAI genes were deleted except *cagX* and *cagY*, which are transcribed as an operon from a putative promoter located in *cag9*, upstream of *cagX* (28, 29). This mutant, designated *cagXY*, is shown schematically in Fig 3 compared to J166 WT and Δ*cagY.* J166 *cagXY* expresses CagY on the bacterial surface (Fig 5A,C) but fails to induce a robust IL-8 response in AGS cells due to the lack of a T4SS (Fig S5). In the flow channel α_5_β_1_ integrin binding assay, J166 *cagXY* binds at a level similar to J166 WT (Fig 5E). To exclude a role for CagX, we deleted all *cag*PAI genes and stitched *cagY* directly to the promoter in *cag9*, creating J166 *cagY*. Similar to J166 *cagXY*, J166 *cagY* fails to induce IL-8 (Fig S5), but expresses CagY and binds to α_5_β_1_ integrin similarly to J166 WT (Fig 5B,D and F). Together these results suggest that in this assay, *H. pylori* binds to α_5_β_1_ integrin predominantly via a CagY-dependent mechanism, but independently of T4SS pilus formation. This conclusion is also supported by the observation that integrin binding in J166 Δ*cagI/L* and Δ*cagE*, which do not form a T4SS pilus (27), is similar to WT (Fig 2D).

**Fig 4:**
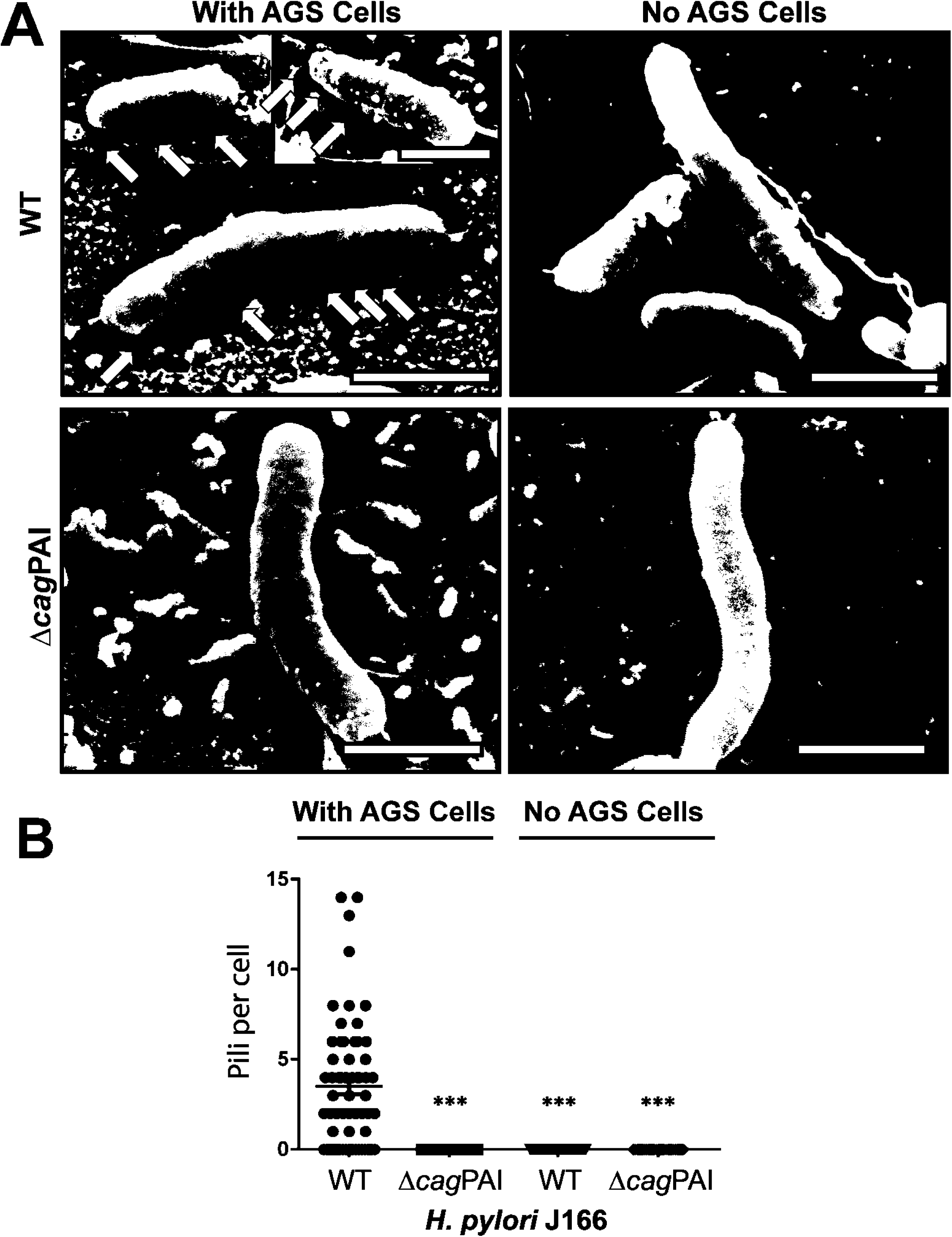
Field emission scanning electron microscopy (FEG-SEM) of *H. pylori* demonstrates that host cell contact is required for T4SS pilus formation. (A) FEG-SEM of *H. pylori* J166 WT and Δ*cag*PAI cultured with or without AGS cells. Pili are indicated by white arrows. Scale bar, 1 µm. (B) Enumeration of pili per bacterial cell. ***P<0.001, compared with WT.

**Fig 5:**
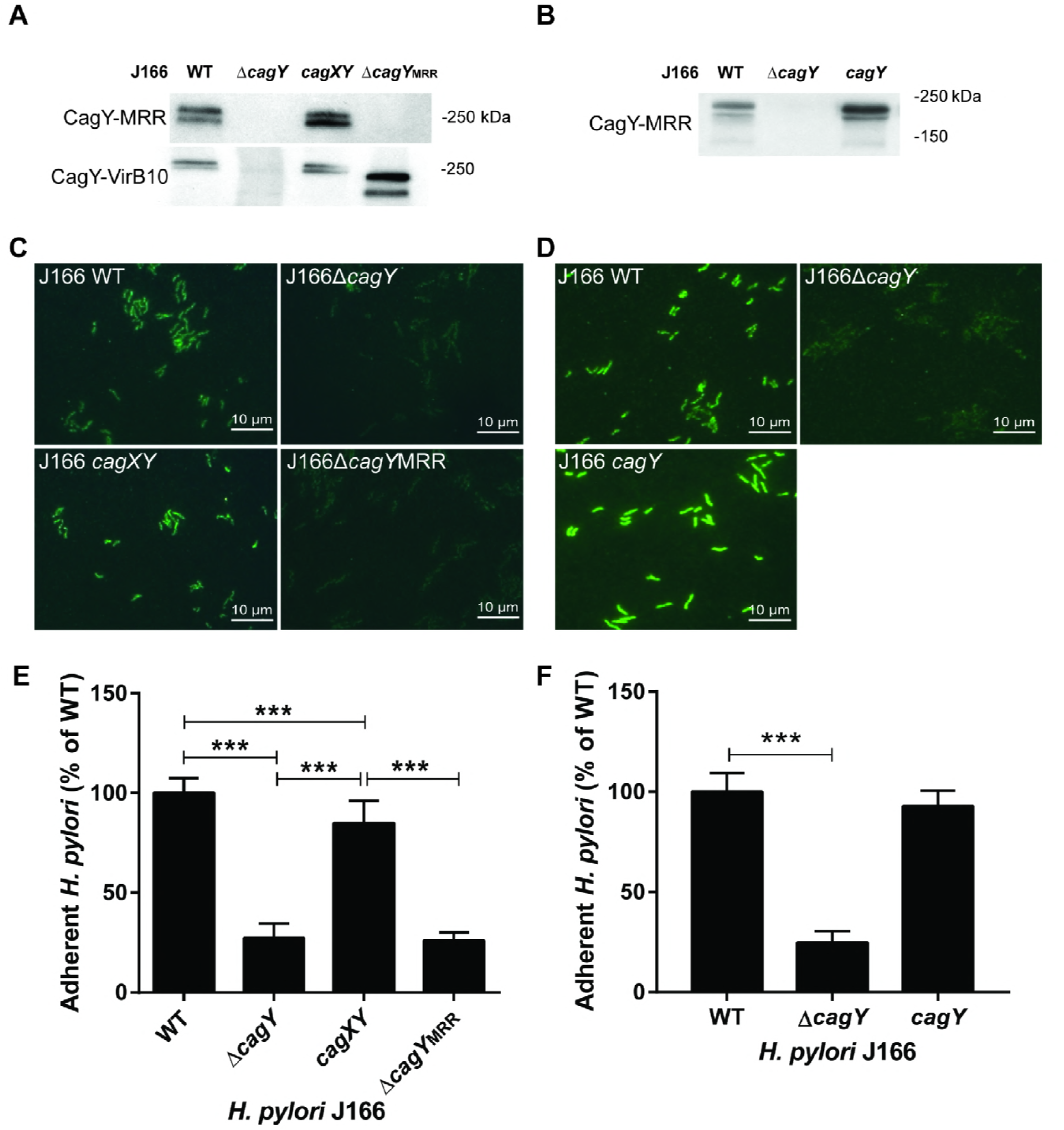
The CagY middle repeat region (MRR), but not the T4SS pilus, is required to bind α_5_β_1_ integrin in a host cell-free system and is expressed on the bacterial surface. (A and B) Immunoblot detection of the CagY MRR and VirB10 region in bacterial lysates. (C and D) Immunofluorescent detection of the CagY MRR on the surface of non-permeabilized *H. pylori*. (E and F) Flow channel competitive adherence to α_5_β_1_ integrin. Results are expressed as the ratio of deletion mutant to WT adherence, and represent the mean (± SEM) of at least 3 independent experiments. ***P<0.001.

### The CagY MRR is necessary for integrin binding

The topography of CagY in the bacterial cell is poorly understood. Proteomic studies suggest that it may be located in the cytoplasmic membrane, or perhaps span the inner and outer membranes [18], similar to what has been demonstrated for the *Escherichia coli* VirB10 (30). However, CagY is much larger than other VirB10 orthologs, and includes two membrane spanning domains that flank the MRR, which previous studies suggested may be localized to the bacterial surface (31). Surface localization is also apparent in deletion mutants of *cagA*, *cagI/L* and *cagE* (Fig S3B), and in J166 *cagXY* (Fig 5A and S3C) and J166 *cagY* (Fig 5D), which do not make a T4SS pilus. We next constructed an unmarked in-frame deletion of the MRR (designated J166Δ*cagY*_MRR_), which is shown schematically in Fig 3. J166Δ*cagY*_MRR_ does not induce IL-8 (Fig S5) or bind α_5_β_1_ integrin in the flow channel (Fig 5E), and, as expected, shows no surface localization of CagY using antibody directed to the MRR (Fig 5C). However, J166Δ*cagY*_MRR_ has an in-frame deletion and produces CagY that can be detected with antibody to the VirB10 portion of CagY (Fig 5A). Together, these results suggest that the *H. pylori* CagY MRR is expressed on the bacterial surface, is required for the binding of α_5_β_1_ integrin in a T4SS-independent manner, and is essential for T4SS function.

### Variation in the motif structure of the CagY MRR alters binding to α_5_β_1_ integrin and T4SS function

We previously demonstrated, using mouse and non-human primate models, that recombination in the *cagY* MRR regulates T4SS function (21, 23), though the mechanism is unknown. Since we have now shown that the MRR is also required to bind α_5_β_1_ integrin in the flow channel, we hypothesized that recombination in *cagY* modulates T4SS function by altering the efficiency of *H. pylori* adhesion to α_5_β_1_ integrin. To test this hypothesis, we compared IL-8 induction to integrin adhesion, using three groups of *H. pylori* strains, each with several isogenic variants bearing unique *cagY* alleles that were previously documented to confer changes in IL-8 induction. First, we examined four isogenic *H. pylori* J166 strains bearing different *cagY* alleles, which arose naturally during infection of mice and were transformed into the WT parent strain (21). All four strains express an unmarked CagY that differs only in the motif structure of the MRR (Fig 6A). Two of the strains induce IL-8 and translocate CagA similarly to WT J166, and two have a non-functional T4SS (21). Consistent with our hypothesis, changes in the J166 CagY MRR that reduced IL-8 also showed a marked and commensurate reduction in adhesion to α_5_β_1_ integrin (Fig 6B). Parallel experiments with isogenic strains of *H. pylori* PMSS1 bearing a unique CagY MRR that altered T4SS function (23) showed similar results (Figs 6C and 6D). Finally, we examined the relationship between induction of IL-8 and integrin binding in paired clonal *H. pylori* isolates recovered from a human patient over a period of 7.4 years (KUS13A and KUS13B), and which differed in the CagY MRR and in T4SS function (23). Again we found that MRR-dependent adhesion of each *H. pylori* isolate to α_5_β_1_ integrin was in most cases commensurate with the level of IL-8 induction (Fig 6E and 6F). Together these results suggest that recombination in *cagY* modulates T4SS function by altering *H. pylori* attachment to α_5_β_1_ integrin.

**Fig 6:**
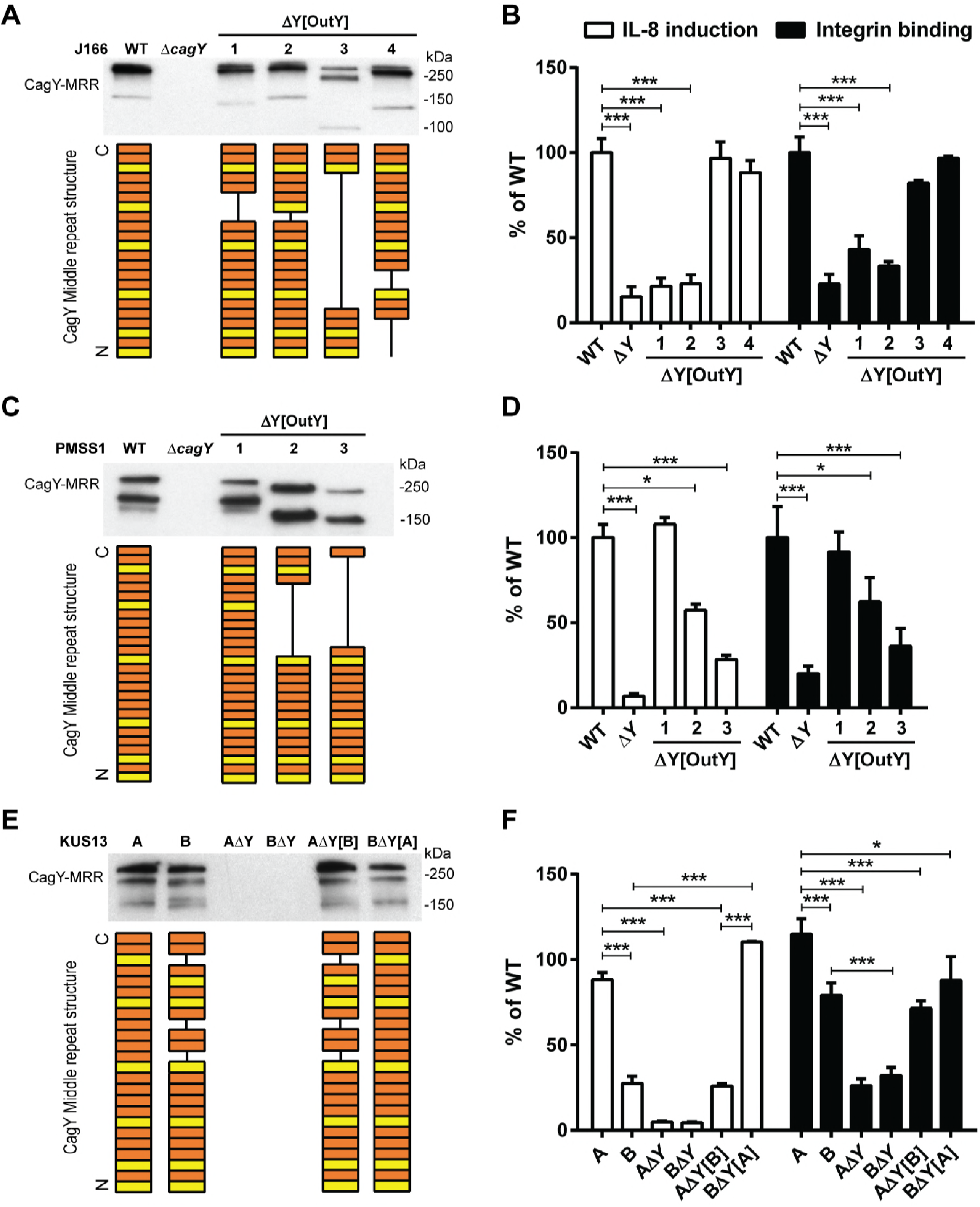
Variation in the amino acid motif structure of the CagY MRR regulates T4SS function by altering α_5_β_1_ integrin binding. Western blot detection of the CagY MRR in whole cell bacterial lysates of *H. pylori* J166 (A) or PMSS1 (C) isogenic strains, each bearing unique *cagY* alleles, or their Δ*cagY* deletion mutants. The corresponding amino acid structure of the MRR is shown schematically as a series of A (orange) or B (yellow) motifs, each 31-39 residues, based on DNA sequence analysis as described previously (61). IL-8 induction (white bars) and integrin adhesion (black bars) relative to WT are shown for *H. pylori* J166 (B) and PMSS1 (D) that correspond to strains shown in panels A and C, respectively. (E) Western blot detection and schematic of the CagY MRR (derived as in panels A and C) from KUS13A, KUS13B, and isogenic variants in which *cagY* was deleted (Δ*cagY*) or replaced with that from the variant strain (i.e., KUS13A with *cagY_13_*_B_ or KUS13B with *cagY_13_*_A_). (F) IL-8 induction (white bars) and integrin adhesion (black bars) relative to WT for the strains shown in panel E. Quantitative results represent the mean (± SEM) of at least 3 independent experiments. *P<0.05, ***P<0.001 for comparison of WT to isogenic *cagY* deletion and variants. Results for IL-8 are adapted from references 20 and 22.

### Variant CagY amino acid motifs that differ in integrin binding and T4SS function are expressed on the bacterial surface

Recombination of *cagY* could modulate integrin binding by changing its amino acid motif structure, but it might also change its level of expression or surface localization. Although in some strains the level of CagY expression appears decreased (e.g. Fig 6A, strain 3), this likely reflects a marked reduction in size of the MRR and reduced antibody recognition. We detected no relationship between MRR expression on western blot and either *H. pylori* adhesion to integrin or induction of IL-8 (Fig 6). CagY MRR was expressed on the bacterial surface in isogenic *H. pylori* PMSS1 strains that differed only in their MRR, and also showed no relationship to T4SS function or integrin binding (Fig 7A). Analysis of fluorescence intensity normalized to DAPI staining demonstrated quantitatively that CagY was expressed on the bacterial surface at similar levels, with no detection in the negative control (Fig 7B). Quantitation of expression on the bacterial surface of isogenic *cag*PAI mutants of J166 similarly showed no relationship to T4SS function or integrin binding, though all MRR variants showed reduced expression (Fig S3D), perhaps related to the reduction in number of MRR motifs. These results suggest that changes in the motif structure of CagY on the bacterial surface modulate T4SS function by altering bacterial adhesion to α_5_β_1_ integrin, rather than altering surface presentation of CagY.

**Fig 7:**
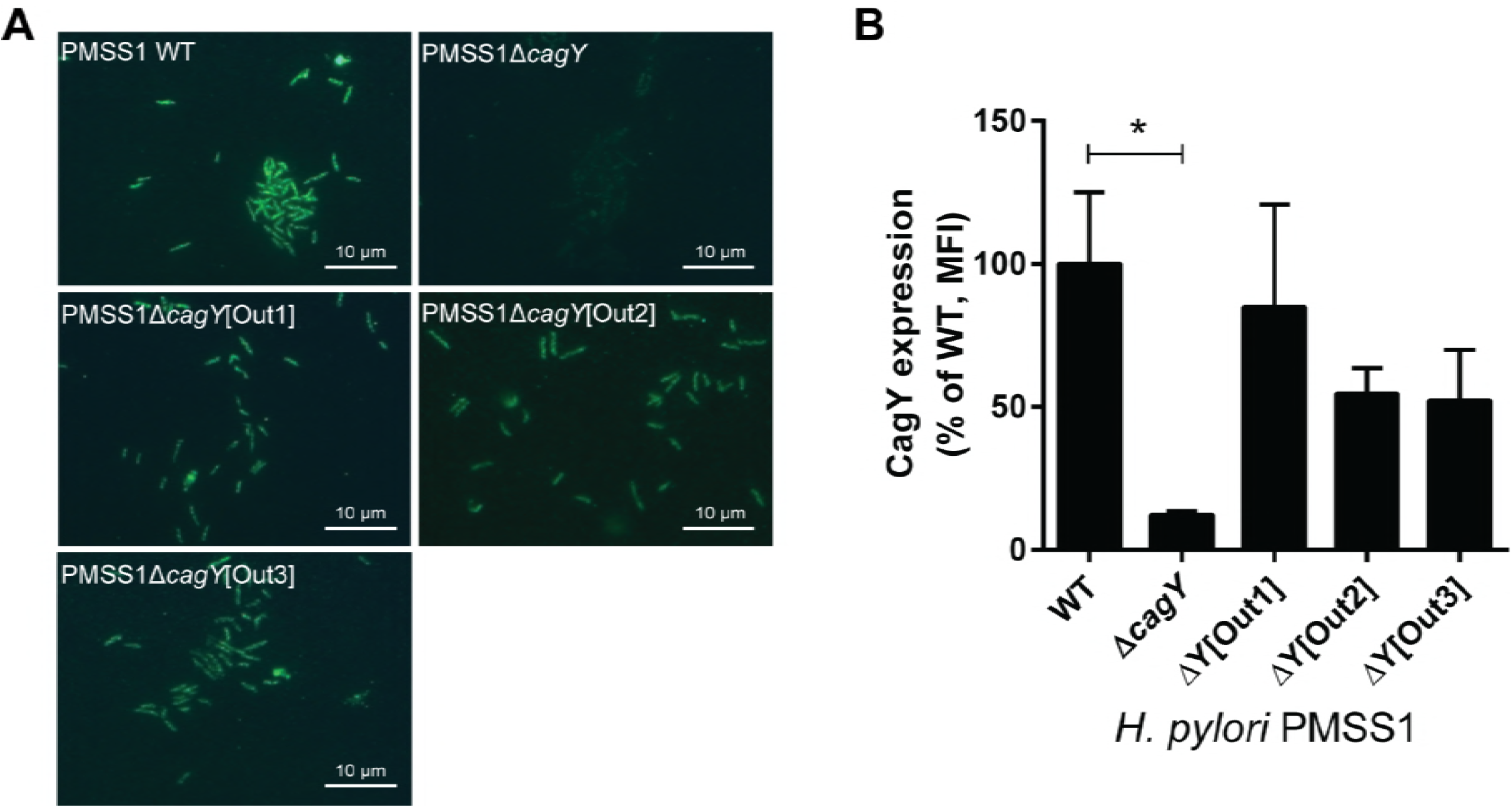
Recombination in *cagY* does not change its surface expression. (A) Immunofluorescent detection of the CagY MRR on the surface of non-permeabilized *H. pylori* PMSS1 with distinct *cagY* alleles from mouse output strains. (B) Quantification of CagY MRR mean fluorescence intensity (MFI) normalized to DAPI. Results are expressed as the percentage ratio of deletion mutant to WT, and represent the mean (± SEM) of 3 independent experiments. *P<0.05, compared to WT.

## DISCUSSION

*H. pylori* persistence in the gastric mucosa is often attributed to evasion of the innate and adaptive immune response, including antimicrobial peptides (32), toll like receptor signaling (33, 34), and T cell proliferation (35, 36), as well as promotion of a regulatory T cell response (37). However, the very presence in most strains of the *cag*PAI, which promotes the host immune response (38, 39), and the uniform occurrence of gastritis in infected patients, suggest the possibility that the host inflammatory response may at the same time actually promote *H. pylori* colonization, a concept that has recently been elegantly demonstrated for several enteric pathogens (40). This is supported by observations of functional antagonism between some *H. pylori* virulence factors such as the CagA oncoprotein and the VacA cytotoxin (41, 42), and by recent evidence that CagA-dependent inflammation may be important for acquisition of essential nutrients such as iron (43, 44) and zinc (45). This more nuanced view of the relationship between *H. pylori* and the host immune response suggests that the overarching strategy used by *H. pylori* to persist in the stomach might be better characterized as immune regulation rather than simply immune evasion.

CagY is an essential component of the *H. pylori* T4SS that may be well-suited to serve this immune regulatory function. The *cagY* gene has in its middle repeat region (MRR) a series of direct DNA repeats that *in silico* predict in-frame recombination events. Recombination in the *cagY* MRR is in fact common, since variants can be readily detected *in vitro*, though it remains possible that the frequency is increased in response to unknown host signals. We previously showed that *cagY* recombination *in vivo* yields a library of insertions and deletions in the MRR, which maintain CagY expression but frequently alter T4SS function (21). CagY-dependent modulation of T4SS function is graded—more like a rheostat than a switch—and can yield variants that confer both gain and loss of function in vivo (21, 23). Adoptive transfer and knockout mouse experiments demonstrate that development of variant *cagY* alleles requires a CD4+ T cell- and IFNγ-dependent immune response (23). Thus, *cagY* recombination can modulate T4SS function and may be a bacterial strategy to both up- and down-regulate the host immune response to promote persistent infection.

Here we addressed the mechanism by which recombination in the MRR alters T4SS function. Since CagY is a ligand for α_5_β_1_ integrin, which is essential for T4SS function, we hypothesized that changes in the amino acid motif structure from recombination in the MRR might alter integrin binding and modulate T4SS function. Analysis of whole bacterial cells in a microfluidic assay demonstrated CagY-dependent and integrin conformation-specific binding to α_5_β_1_, which correlated closely with T4SS function in isogenic variants that differed only in the MRR region of CagY. This binding was independent of the T4SS pilus, which was not formed under these cell-free conditions, though the MRR was expressed on the bacterial surface as described previously (31). Moreover, we could detect the MRR on the bacterial surface even when the entire *cag*PAI was deleted except for *cagY* and the upstream promoter. Binding to α_5_β_1_ integrin was not dependent upon CagA, CagE, or CagL, which was originally identified as the ligand for α_5_β_1_ integrin (11). While CagL is clearly essential for T4SS function, more recent studies have suggested that it binds α_V_β_6_ and α_V_β_8_ integrin and not α_5_β_1_ (17). The results from yeast two-hybrid studies (12) also identified CagI as a β_1_ integrin binding partner, which we could not confirm in whole bacterial cells.

Previous studies have found that the VirB10 ortholog at the C-terminus of CagY bound to α_5_β_1_ integrin, but not the MRR region. However, these studies examined protein-protein interactions by yeast two-hybrid and immunoprecipitation or by surface plasmon resonance (12, 46), which may not reflect binding in a whole bacterial cell. Since the isogenic *cagY* variants examined here differed only in the MRR, and deletion of the MRR eliminated α_5_β_1_ integrin binding, our results suggest that the *H. pylori* MRR is required for binding to α_5_β_1_ integrin in an intact bacterial cell. However, we have not directly examined MRR binding to α_5_β_1_ integrin, so the MRR may not itself be an integrin ligand, but instead may modulate binding of the VirB10 domain of CagY. We have been unable to demonstrate CagY-dependent adherence to α_5_β_1_ integrin on AGS gastric epithelial cells in our microfluidic assay (data not shown), which may reflect the multiple binding partners, including *cag*PAI components, as well as HopQ, BabA, SabA, and other outer membrane adhesins (47, 48). Others have also found no difference in binding to AGS cells between WT and Δ*cag*PAI (12).

The topology of CagY in the bacterial membrane, and the accessibility to the α_5_β_1_ integrin, also remain areas of uncertainty. Integrins are generally found in the basolateral compartment, which would not normally be accessible to *H. pylori* on the apical cell surface. However, *H. pylori* binds preferentially at tight junctions in cell culture and in gastric tissue, leading to disruption of the integrity of the epithelial layer (49). Moreover, recent studies suggest that *H. pylori* HtrA, an essential serine protease, cleaves occludin, claudin-8, and E-cadherin, which opens cell-cell junctions and may explain how *H. pylori* could bind integrins *in vivo* (50-52). *H. pylori* binding to CEACAMs (53, 54) or other yet identified cell receptors may also induce redistribution of integrins from the basolateral to the apical cell surface, making them accessible to CagY. It also remains unclear precisely how CagY is localized in the bacterial cell membrane. Elegant cryo-electron microscopy studies have demonstrated that the VirB10 orthologue in the *Escherichia coli* plasmid conjugation T4SS forms part of a core complex that spans the inner and outer bacterial membranes (30). However, the topology in *H. pylori* appears different, as recent electron microscopy studies suggest that the core complex is much larger than that in *E. coli*, and is composed of 5 proteins (rather than 3), including CagX, CagY, CagM, CagT, and Cag3 (25).

In conclusion, these studies demonstrate that CagY modulates attachment to α_5_β_1_ integrin independently of the T4SS pilus in a manner that depends on the MRR motif structure. It is tempting to speculate that CagY-mediated alteration in integrin binding is also mechanistically linked to T4SS function, since they are strongly correlated (Figure 6). For example, surface expression of an integrin-binding motif may promote intimate epithelial cell contact, which in turn serves as a nucleation signal to promote expression of the T4SS pilus, further enhancing integrin binding and injection of effector molecules. Such a scenario might entail MRR-dependent integrin signaling, including activation of focal adhesion kinase (FAK) and the Src family kinase, though others have shown that only the extracellular domains of the β_1_ integrin are important for CagA translocation (12). On the other hand, it is logically possible that changes in the MRR affect integrin binding and T4SS function independently. Though the details remain to be elucidated, we hypothesize that CagY-dependent binding to α_5_β_1_ integrin serves as a molecular rheostat that “tunes” the optimal balance between the competing pressures of gastric inflammation, which serves a metabolic function for the bacterium on the one hand, but comes at a cost of exposure to immune pressure, decreased bacterial load, and decreased possibility of transmission to a new host.

## MATERIALS AND METHODS

### Construction and culture of *E. coli* expressing InvA

Plasmid pRI253 (kindly provided by Ralph Isberg, Tufts University, Boston, MA) contains the *invA* gene from *Yersinia pseudotuberculosis* under the control of a phage T7 RNA polymerase promoter (55). To create a negative control, the *invA* gene was cut out from the plasmid using restriction enzymes EcoRI and HindIII. The recircularized plasmid pRI253Δ*invA* and the original plasmid pRI253 were transformed into competent *E. coli* BL21 (Invitrogen) according to the manufacturers’ instructions. *E. coli* strains were cultured overnight at 37°C in Luria Bertani (LB) broth supplemented with 5 mg/liter carbenicillin. Overnight cultures were diluted to an OD_600_ of 0.05, cultured for an additional 2-3 h, followed by addition of 0.5 mM Isopropyl β-D-1-thiogalactopyranoside (IPTG) and another 2 h of incubation to induce InvA expression.

### *H. pylori* strains and culture conditions

Wild type *H. pylori* strains were cultured on Brucella agar or in Brucella broth (BBL/Becton Dickinson, Sparks, MD) supplemented with 5% heat-inactivated newborn calf serum (Invitrogen, Carlsbad, CA) and antibiotics (trimethoprim, 5 mg/liter; vancomycin, 10 mg/liter; polymyxin B, 2.5 IU/liter, amphotericin B, 2.5 mg/liter). *H. pylori* mutant strains were cultured as for wild type, but with the addition of kanamycin (25 mg/liter), chloramphenicol (5 mg/liter), or streptomycin (10 mg/liter) as appropriate (all antibiotics from Sigma). *H. pylori* liquid cultures were grown overnight to an optical density at 600 nm (OD_600_) of approximately 0.3 to 0.4. All *H. pylori* cultures were grown at 37°C under microaerophilic conditions generated by a fixed 5% O_2_ concentration (Anoxomat, Advanced Instruments, Norwood, MA). A complete list of strains is shown in Table 1.

**Table 1.**
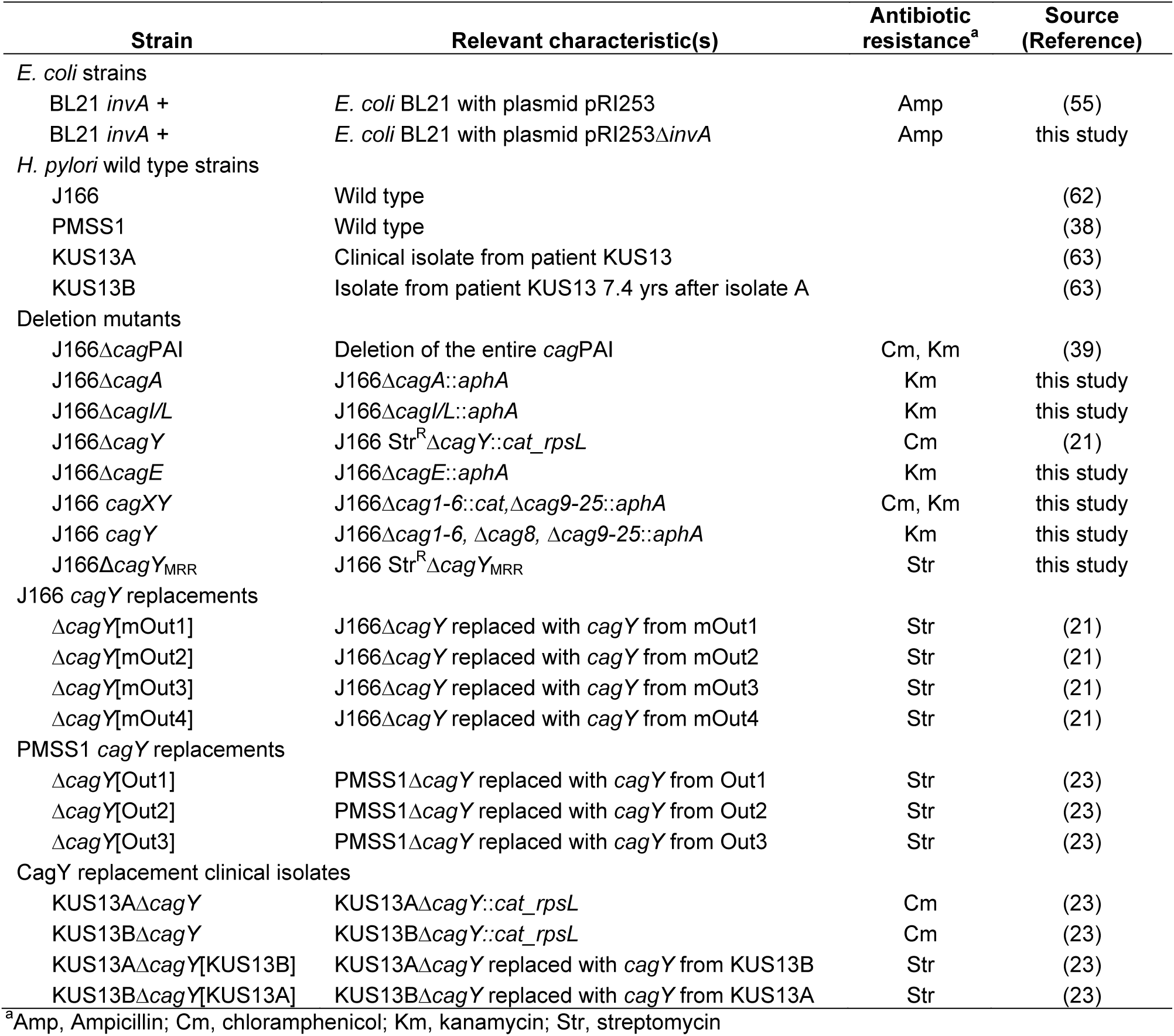
*H. pylori* strains

| Strain | Relevant characteristic(s) | Antibiotic resistance<sup>a</sup> | Source (Reference) |
| --- | --- | --- | --- |
| *E. coli* strains |  |  |  |
| BL21 *invA* + | *E. coli* BL21 with plasmid pRI253 | Amp | (55) |
| BL21 *invA* + | *E. coli* BL21 with plasmid pRI253Δ*invA* | Amp | this study |
| *H. pylori* wild type strains |  |  |  |
| J166 | Wild type |  | (62) |
| PMSS1 | Wild type |  | (38) |
| KUS13A | Clinical isolate from patient KUS13 |  | (63) |
| KUS13B | Isolate from patient KUS13 7.4 yrs after isolate A |  | (63) |
| Deletion mutants |  |  |  |
| J166Δ*cagPAI* | Deletion of the entire *cagPAI* | Cm, Km | (39) |
| J166Δ*cagA* | J166Δ*cagA::aphA* | Km | this study |
| J166Δ*cagI/L* | J166Δ*cagI/L::aphA* | Km | this study |
| J166Δ*cagY* | J166 Str<sup>R</sup>Δ*cagY::cat_rpsL* | Cm | (21) |
| J166Δ*cagE* | J166Δ*cagE::aphA* | Km | this study |
| J166 *cagXY* | J166Δ*cag1-6::cat,Δcag9-25::aphA* | Cm, Km | this study |
| J166 *cagY* | J166Δ*cag1-6, Δcag8, Δcag9-25::aphA* | Km | this study |
| J166Δ*cagY*<sub>MRR</sub> | J166 Str<sup>R</sup>Δ*cagY*<sub>MRR</sub> | Str | this study |
| J166 *cagY* replacements |  |  |  |
| Δ*cagY*[mOut1] | J166Δ*cagY* replaced with *cagY* from mOut1 | Str | (21) |
| Δ*cagY*[mOut2] | J166Δ*cagY* replaced with *cagY* from mOut2 | Str | (21) |
| Δ*cagY*[mOut3] | J166Δ*cagY* replaced with *cagY* from mOut3 | Str | (21) |
| Δ*cagY*[mOut4] | J166Δ*cagY* replaced with *cagY* from mOut4 | Str | (21) |
| PMSS1 *cagY* replacements |  |  |  |
| Δ*cagY*[Out1] | PMSS1Δ*cagY* replaced with *cagY* from Out1 | Str | (23) |
| Δ*cagY*[Out2] | PMSS1Δ*cagY* replaced with *cagY* from Out2 | Str | (23) |
| Δ*cagY*[Out3] | PMSS1Δ*cagY* replaced with *cagY* from Out3 | Str | (23) |
| CagY replacement clinical isolates |  |  |  |
| KUS13AΔ*cagY* | KUS13AΔ*cagY::cat_rpsL* | Cm | (23) |
| KUS13BΔ*cagY* | KUS13BΔ*cagY::cat_rpsL* | Cm | (23) |
| KUS13AΔ*cagY*[KUS13B] | KUS13AΔ*cagY* replaced with *cagY* from KUS13B | Str | (23) |
| KUS13BΔ*cagY*[KUS13A] | KUS13BΔ*cagY* replaced with *cagY* from KUS13A | Str | (23) |

<sup>a</sup>Amp, Ampicillin; Cm, chloramphenicol; Km, kanamycin; Str, streptomycin

### Construction of *H. pylori* mutants

Six mutants in *H. pylori* J166 were constructed (Table 1). For J166Δ*cagA*, J166Δ*cagI/L* and J166Δ*cagE*, DNA fragments upstream and downstream of the respective gene deletion were PCR amplified using primers (Table S1) with restriction sites that permitted ligation to a kanamycin resistance gene (*aphA*), and insertion into the multiple cloning site of pBluescript (Stratagene, La Jolla, CA). The resulting plasmid was transformed into *E. coli* TOP10 (Invitrogen) according to the manufacturers’ instructions, and transformants were grown overnight on Luria-Bertani (LB) plates containing kanamycin. Resistant colonies were inoculated in selective LB broth and plasmids from the resulting culture were purified with a QIAprep^®^ Spin Miniprep kit (Qiagen). Plasmids were sequenced and digested with appropriate enzymes for verification of correct construction prior to natural transformation of *H. pylori* with kanamycin selection. J166 *cagXY* was created in a similar fashion, but in two steps, first deleting *cag1-6* with a chloramphenicol resistance cassette (*cat*) and selection on chloramphenicol, and then deleting *cag9-25* with a kanamycin cassette, leaving only *cagX*, *cagY* (and its promoter), and *cagA*, which in strain J166 is not on the *cag*PAI (56). J166 *cagY* was made in a series of 3 steps. First an unmarked deletion of *cag1-6* was constructed using contraselection. The region was replaced by a *cat*-*rpsL* casette, resulting in streptomycin sensitive (*rpsL* encodes dominant streptomycin sensitivity) and chloramphenicol resistant transformants. Then upstream and downstream fragments were stitched together and the PCR product was used to replace the cassette, leaving an unmarked deletion. Next, *cag9-25* were deleted as in the *cagXY* construct, and replaced with a kanamycin cassette. Finally, contraselection was again used to excise the *cagX* gene, bringing 313 bp upstream of *cagX* (putatively containing its promoter) immediately upstream of *cagY*.

J166Δ*cagY*_MRR_, with an in-frame markerless deletion of the MRR, was constructed using modifications of contraselection described previously (21). Briefly, the MRR was first replaced by insertion of the *cat*-*rpsL* cassette in streptomycin resistant *H. pylori* J166. Fragments upstream and downstream of the MRR were then each amplified with overlapping primers that permitted stitching of the two products in a second PCR reaction. The stitched product was ligated into pBluescript and used in a second transformation reaction to replace *cat*-*rpsL*, with selection on streptomycin. All *H. pylori* deletion mutants were sequence verified to confirm the correct construction.

### Microfluidic adhesion assay

Microfluidic adhesion assays were assembled as previously described (57). In brief, 25 mm diameter, #1.5 glass coverslips were piranha etched to remove organic molecules and treated with 1% 3-aminopropyltriethoxysilane to add aminosilane groups. Recombinant human α_4_β_1_, α_5_β_1_ or α_L_β_2_ integrin (R&D Systems, Minneapolis, MN) was adsorbed at 10 mg/liter concentration overnight at 4°C resulting in approximately 2000 sites/μm^2^. Coverslips were then washed and blocked with Hank’s balanced salt solution with 0.1% human serum albumin. Where indicated, the blocking solution was supplemented with 5 mg/liter of anti-integrin β_1_ blocking antibody P5D2 (Abcam, San Francisco, CA), low affinity locking anti-integrin β_1_ antibody SG19, high affinity locking anti-integrin β_1_ antibody TS2/16 (both from Biolegend, San Diego, CA), isotype control antibody B11/6 (Abcam, San Francisco, CA), or 2 mM MnCl_2_ (Mn^2+^) to activate integrin. Custom multi-channel microfluidic device (57) was vacuum sealed, outlets were attached to Exigo pumps to provide the negative pressure necessary to induce shear, and inlet reservoirs were loaded with *E. coli* or *H. pylori*. Prior to loading, liquid cultured bacteria were stained at an OD_600_ of 0.8 with 2% Vybrant DiI, DiD or DiO Cell-Labeling Solution (Grand Island, NY) in Brucella broth for 20 min at 37°C in the dark. Stained bacteria were washed twice with PBS and then resuspended in Brucella broth to the desired final OD_600_. Competitive binding assays were performed by mixing differently labeled WT and mutant bacteria at an OD_600_ of 0.4 (total OD_600_ 0.8). Shear was induced at 1 dyne/cm^2^ for 3 minutes followed by a 3 minute period of no shear incubation to allow attachment. Then shear was increased to 1 dyne/cm^2^ and 10 second videos were taken along the centerline of the channel in four field of views using an inverted TIRF research microscope (Nikon) equipped with a 60X numerical aperture 1.5 immersion TIRF objective and a 120 W arc lamp to capture epi-fluorescence images with the appropriate filter sets (488 nm for DiO, 510 nm for DiD and 543 nm for DiI). Images were captured using a 16-bit digital complementary metal oxide semiconductor Zyla camera (Andor, Belfast BT12 7AL, UK) connected to a PC (Dell) with NIS Elements imaging software (Nikon, Melville, NY). Images were collected with 2×2 binning at a resolution of 1024 x 1024 at a rate of 2 frames per second. Adherent bacteria were identified by the presence of fluorescence, which was cross-checked with an overlaid brightfield image to eliminate fluorescent noise. Small numbers of bacteria that were unstained (typically ~10%) were not counted. Bacteria that remained stationary or tethered after 10 seconds were counted visually in 3 fields of view, and the results were averaged for each biological replicate. To assess reliability, two observers (one blinded) independently scored adherent bacteria at 488 nm and 543 nm in 9 fields of view that contained competitive binding assays (WT and mutant). Mean similarity for the 18 observations was 0.94, which was calculated as 1-[|O_1_ - O_2_|/ ½(O_1_ + O_2_)], where O_1_ and O_2_ are the independent scores for the two observers, and a value of 1.0 indicates perfect agreement. Data on integrin binding are representative of at least three biological replicates, which in most cases examined three fields of view in duplicate technical replicates.

### Sequencing of *cagY*

The DNA sequences of *cagY* from *H. pylori* PMSS1 and KUS13A and B were determined using single molecule real-time sequencing (Pacific Biosciences, Menlo Park, CA). Briefly, *cagY* was amplified as previously described (21) and purified PCR products were submitted to the DNA Technologies Core at the UC Davis Genome Center. The amplicons were sequenced using a PacBio RSII sequencer, with P6C4 chemistry. Data were analyzed using PacBio’s SMRTportal Analysis 2.3.0. Sequences were deposited in GenBank and are available under accession numbers KY613376 - KY613380. *cagY* sequences of *H. pylori* J166 were previously published (21).

### Assessment of protein expression by fluorescence microscopy

Liquid cultures of *H. pylori* or IPTG-induced *E. coli* strains were centrifuged (3,000 x g, 3 min) and resuspended in blocking buffer (PBS with 1% bovine serum albumin and 0.05% Tween 20) at an OD_600_ of 0.4. Each culture was spotted on to two microscope slides using cytofunnels in a cytospin at 1000 rpm for 15 min. Air dried slides were incubated for 1 h with blocking buffer in a humid chamber followed by 1 h incubation with anti-*H. pylori* CagY MRR antibody (31) diluted 1:1000 or anti-Yersinia invasin antibody (58) diluted 1:5000 in blocking buffer. Slides were washed 3 times with PBS and incubated for 1 h in the dark with Alexa Fluor 488 goat anti-rabbit IgG (R37116, Life Technologies) diluted 1:10 in blocking buffer. After further washing, the slides were mounted with FluoroShield with DAPI (Sigma). The slides were stored in the dark and imaged the next day. Photos of all slides were captured with the same exposure time for each antibody and DAPI. Fluorescence intensity was analyzed with the ImageJ software, normalizing the total CagY fluorescence at a given threshold determined by the positive WT sample to the area of the DAPI fluorescence of the bacterial particles.

### Immunoblots

Expression of *E. coli* invasin and *H. pylori* CagY MRR were detected by electrophoresis of lysates of liquid cultured bacteria as described previously (21), using polyclonal rabbit antisera to invasin (1:15,000) or CagY MRR (1:10,000) as primary antibodies. Detection of CagY expression in Δ*cagY*_MRR_, an in-frame deletion of the MRR, was performed using antiserum from rabbits immunized with the VirB10 portion at the C-terminus of CagY (1:1,000). To generate the antiserum, DNA encoding the C-terminus of *H. pylori* J166 CagY was PCR amplified (Table S1), cloned into pGEX-4T-3 vector and transformed into *E. coli* BL21 (both GE Healthcare). Expression of the GST-fusion protein and preparation of cell extracts was performed according to manufacturer’s instructions. The GST-fusion protein was bound to Glutathione Sepharose 4B (GE Healthcare) in a column and the GST was cleaved off by thrombin. The eluate was run on a SDS-PAGE, the purified CagY C-terminus protein was cut out from the gel and was used to generate rabbit antisera according to standard protocols (Antibodies, Inc, Davis, CA).

### IL-8 ELISA

IL-8 was measured essentially as described previously (59). Human AGS gastric adenocarcinoma cells (ATCC, Manassas, VA) were grown in RPMI 1640 supplemented with 10% fetal bovine serum, 100 units/mL penicillin and 100 μg/mL streptomycin at 5% CO_2_, 37°C. All antibiotics were excluded from the growth media 24 h prior to *H. pylori* co-culture. Approximately 5×10^5^ human AGS gastric adenocarcinoma cells were seeded in six well plates with 1.8 ml RPMI/10% fetal bovine serum, incubated overnight, and then co-cultured with bacteria diluted in 200 μl Brucella broth to give an MOI of 100:1. Brucella broth with no bacteria served as a baseline control. Supernatants were harvested after 20-22 hours of culture (37°C, 5% CO2), stored at -80°C, and then diluted 1:8 prior to IL-8 assay by ELISA (Invitrogen, Camarillo, CA) performed according to the manufacturer’s protocol. WT *H. pylori* J166 or PMSS1 and its isogenic *cagY* deletion were included on every plate as positive and negative controls, respectively. IL-8 values were normalized to WT *H. pylori* determined concurrently.

### High resolution field-emission gun scanning electron microscopy analyses

Bacteria were cultured alone or with AGS cells for 4 hrs at an MOI of 100:1. Bacteria were prepared for scanning electron microscopy as previously described (27, 60). Briefly, samples were cultured on poly-L-lysine coated glass coverslips, and fixed for 1 hour with 2.0% paraformaldehyde/2.5% gluteraldehyde in 0.05 M sodium cacodylate buffer. Cells were washed three times in 0.05 M sodium cacodylate buffer before secondary fixation with 0.1% osmium tetroxide for 15 minutes. Three additional 0.05 M sodium cacodylate buffer washes were performed before subjecting the samples to sequential ethanol dehydration. Cells were dried at the critical point and carbon-coated before imaging with an FEI Quanta 250 FEG-SEM. Pili were enumerated in a blinded fashion using ImageJ software.

### Statistical analysis

Data are reported as mean ± SEM. Multiple groups were compared using ANOVA, with Tukey’s or Bonferroni’s post hoc test, or with Dunnett’s post hoc test when compared only to WT. Two group comparisons were performed using Student’s *t* test. All analyses were carried out using GraphPad Prism 5.01 for Windows (GraphPad Software, San Diego, CA). A P-value <0.05 was considered statistically significant.

## ACKNOWLEDGEMENTS

We thank Jordan Feeney, UC Davis, CA, for drawing assistance with Figure 1A, Ralph Isberg, Tufts University, Boston, MA, for providing pRI253 plasmid and Virginia Miller, University of North Carolina, Chapel Hill, NC, for providing anti-invasin antibody.

## SUPPORTING FIGURE LEGENDS

**S1 Fig: Adherence of invasin expressing *E. coli*.** (A) Immunoblot of *E. coli* with plasmid pRI253 with or without *invA*, untreated or treated with IPTG to induce InvA expression. Full length InvA is around 100 kDa and the lower bands represent degradation products (55, 58). (B) Attachment to α_5_β_1_ integrin in a competitive assay; the two strains of *E. coli* were stained with either DiD or DiO and mixed 1:1 for a total OD_600_ of 0.8. A parallel dye swap was performed to exclude influence of the dye on the integrin attachment.

**S2 Fig. Fluorophore labeling control experiments.** (A) Percent of cells stained with Dil or DiO ([fluorescently stained cells divided by total cells seen on brightfield] x 100) was determined for *H. pylori* J166 WT, Δ*cag*PAI, Δ*cagY*, and isogenic strains expressing functional (Δ*cagY*[Out1]) or non-functional (Δ*cagY*[Out3] *cagY* alleles. (B) *H. pylori* J166 WT and Δ*cag*PAI (each at OD_600_=0.4) were labeled with DiI or DiO, respectively (left), mixed 1:1, and imaged with a 546 nm (DiI) or 488 nm (DiO) light source. Parallel experiments were performed with a dye swap (middle). WT labeled with DiI was also mixed with WT labeled with DiO (right). Binding to α_5_β_1_ integrin was similar whether detected with DiO or Dil fluorescent dyes, and whether in competition with a strain with low or high binding ability. Results for WT measured in a competition setting were also very similar to that when measured individually (compare with Fig 2B). Data represent mean ± SEM of ≥ 3 experiments.

**S3 Fig. Expression of CagY in wild type and isogenic knockouts of *H. pylori* J166.** (A) Immunoblot of CagY in WT *H. pylori* and isogenic knockouts of *cagY*, *cagA*, *cagI/L*, and *cagE*. Immunoblot of UreB is shown as a loading control. (B-D) Quantification of whole cell CagY MRR fluorescence signal normalized to DAPI in non-permeabilized WT *H. pylori* and isogenic knockouts or *cagY* variants. CagY surface expression in strains with MRR motifs that bind α_5_β_1_ integrin and have a functional *cag*PAI (variants 1 and 2) is similar to that in strains that do not bind integrin and do not have a functional *cag*PAI (variants 3 and 4), though all are generally lower than J166 WT. Results are expressed as the percentage ratio of deletion mutant to WT, and represent the mean (± SEM) of 3 independent experiments. (MFI=mean fluorescence intensity) *P<0.05, **P<0.01, ***P<0.001 compared to WT.

**S4 Fig. Field emission scanning electron microscopy (FEG-SEM) of *H. pylori* PMSS1.** (A) FEG-SEM of *H. pylori* PMSS1 WT and Δ*cag*PAI cultured with or without AGS cells. Pili are indicated by white arrows. (B) Enumeration of pili per bacterial cell. ***P<0.001.

**S5 Fig. T4SS function in various *cag*PAI *H. pylori* J166 mutants.** IL-8 induction in AGS cell co-cultured with J166 WT, Δ*cagY*, *cagXY*, *cagY* and Δ*cagY*_MRR_. ***P<0.001.

**S1 Table. Primers used for PCR.**

